# PyDamage: automated ancient damage identification and estimation for contigs in ancient DNA *de novo* assembly

**DOI:** 10.1101/2021.03.24.436838

**Authors:** Maxime Borry, Alexander Hübner, A.B. Rohrlach, Christina Warinner

## Abstract

DNA *de novo* assembly can be used to reconstruct longer stretches of DNA (contigs), including genes and even genomes, from short DNA sequencing reads. Applying this technique to metagenomic data derived from archaeological remains, such as paleofeces and dental calculus, we can investigate past microbiome functional diversity that may be absent or underrepresented in the modern microbiome gene catalogue. However, compared to modern samples, ancient samples are often burdened with environmental contamination, resulting in metagenomic datasets that represent mixtures of ancient and modern DNA. The ability to rapidly and reliably establish the authenticity and integrity of ancient samples is essential for ancient DNA studies, and the ability to distinguish between ancient and modern sequences is particularly important for ancient microbiome studies. Characteristic patterns of ancient DNA damage, namely DNA fragmentation and cytosine deamination (observed as C-to-T transitions) are typically used to authenticate ancient samples and sequences. However, existing tools for inspecting and filtering aDNA damage either compute it at the read level, which leads to high data loss and lower quality when used in combination with de novo assembly, or require manual inspection, which is impractical for ancient assemblies that typically contain tens to hundreds of thousands of contigs. To address these challenges, we designed PyDamage, a robust, automated approach for aDNA damage estimation and authentication of *de novo* assembled aDNA. PyDamage uses a likelihood ratio based approach to discriminate between truly ancient contigs and contigs originating from modern contamination. We test PyDamage on both simulated, and empirical aDNA data from archaeological paleofeces, and we demonstrate its ability to reliably and automatically identify contigs bearing DNA damage characteristic of aDNA. Coupled with aDNA *de novo* assembly, PyDamage opens up new doors to explore functional diversity in ancient metagenomic datasets.

## INTRODUCTION

Ancient DNA (aDNA) is highly fragmented (Orlando et al., 2021; Warinner et al., 2017). Although genomic DNA molecules within a living organism can be millions to hundreds of millions of base pairs (bp) long, postmortem enzymatic and chemical degradation after death quickly reduces DNA to fragment lengths of less than 150 bp, typically with medians less than 75 bp and modes less than 50 bp (Mann et al., 2018; Hansen et al., 2017). Within the field of metagenomics, many approaches require longer stretches of DNA for adequate analysis, a requirement that particularly applies to functional profiling, which often involves *in silico* translation steps (Seemann, 2014). For example, FragGeneScan (Rho et al., 2010), a tool designed for gene prediction from short read data, fails to predict open-reading frames in DNA sequences shorter than 60 bp. If applied directly to highly fragmented ancient metagenomic datasets, such data filtering can introduce biases that interfere with functional analyses when preservation is variable across samples or when comparing ancient samples to modern ones.

Because very short (<100 bp) and ultrashort (<50 bp) DNA molecules pose many downstream analytical challenges, there is a long-standing interest in leveraging the approach of *de novo* assembly to computationally reconstruct longer stretches of DNA for analysis. With de novo assembly, longer contiguous DNA sequences (contigs), and sometimes entire genes or gene clusters, can be reconstructed from individual sequencing reads (Compeau et al., 2011), which can then be optionally binned into metagenome-assembled genomes (MAGs) (Kang et al., 2015). Such contigs are more amenable to functional profiling, and applying this technique to microbial metagenomics datasets derived from archaeological remains, such as paleofeces and dental calculus, has the potential to reveal ancient genes and functional diversity that may be absent or underrepresented in modern microbiomes (Tett et al., 2019; Wibowo et al., 2021; Brealey et al., 2020). However, because ancient samples generally contain a mixture of ancient bacterial DNA and modern bacterial contaminants, it is essential to distinguish, among the thousands of contigs generated by assembly, truly ancient contigs from contigs that may originate from the modern environment, such as the excavation site, storage facility, or other exogenous sources.

In addition to being highly fragmented, aDNA also contains other forms of characteristic molecular decay, namely cytosine deamination (observed as C-to-T transitions in aDNA datasets) (Dabney et al., 2013), which can be measured and quantified to indicate the authenticity of an ancient sample, or even an individual sequence (Hofreiter et al., 2001; Briggs et al., 2007b). However, tools for inspecting and filtering aDNA damage were primarily designed for genomic and not metagenomic applications, and they are largely unsuited or impractical for use in combination with *de novo* assembly. For example, PMDTools (Skoglund et al., 2014) operates at the read level, and when subsequently combined with *de novo* assembly leads to higher data loss and lower overall assembly quality. MapDamage (Ginolhac et al., 2011) and DamageProfiler (Neukamm et al., 2020) are tools that can be applied to assembled contigs, but require manual contig inspection by the user, which is infeasible for *de novo* assemblies yielding tens to hundreds of thousands of contigs. Other tools, such as mapDamage2 (Jónsson et al., 2013), do provide an estimation of damage, but use slower algorithms that do not scale well to the analysis of many thousands of contigs. A faster, automated approach with a better sensitivity for distinguishing truly ancient contigs from modern environmental contigs is needed.

Here, we present PyDamage, a software tool to automate the process of contig damage identification and estimation. PyDamage models aDNA damage from deamination data (C-to-T transitions), and tests for damage significance using a likelihood ratio test to discriminate between truly ancient contigs and contigs originating from modern contaminants. Testing PyDamage on *in silico* simulated data, we show that it is able to accurately distinguish ancient and modern contigs. We then apply PyDamage to de novo assembled DNA from ancient paleofeces from the site of Cueva de los Muertos Chiquitos, Mexico (ca. 1300 BP) and find that the contigs PyDamage identifies as ancient are consistent with taxa known to be members of the human gut microbiome. Among the ancient contigs, PyDamage authenticated multiple functional genes of interest, including a multidrug and bile salt resistance gene cluster from the gut microbe *Treponema succinifacians*, a species that is today only found in societies practicing traditional forms of subsistence. Using PyDamage, *de novo* assembled contigs from aDNA datasets can be rapidly and robustly authenticated for a variety of downstream metagenomics applications.

## MATERIAL AND METHODS

### Simulated sequencing data

In order to evaluate the performance of PyDamage with respect to the GC content of the assembled genome, the sequencing depth along the genome, the amount of observed aDNA damage on the DNA fragments, and the mean length of these DNA fragments, we simulated short-read sequencing data using gargammel (Renaud et al., 2017) varying these four parameters. We chose three microbiome-associated microbial taxa with low (*Methanobrevibacter smithii*, 31%), medium (*Tannerella forsythia*, 47%), and high (*Actinomyces dentalis*, 72%) GC content, following Mann et al. (2018) (Figure 1a). Using three different read length distributions (Figure 1b), we generated short-read sequencing data from each reference genome using gargammel’s *fragSim*. To the resulting short-read sequences we added different amounts of aDNA damage using gargammel’s *deamSim* so that ten levels of damage ranging from 0% to 20% were observed, which were measured as the amount of observed C-to-T substitutions on the terminal base at the 5’ end of the DNA fragments (Figure 1c). Finally, each of these 90 simulated datasets was subsampled to generate nine coverage bins ranging from 1-fold to 500-fold genome coverage by randomly drawing a coverage value from the uniform distribution defining each bin (Figure 1d) and these were aligned to their respective reference genome using BWA *aln* (Li and Durbin, 2009) with the non-default parameters optimized for aDNA *-n 0.01 -o 2 -l 16500* (Meyer et al., 2012).

**Figure 1.**
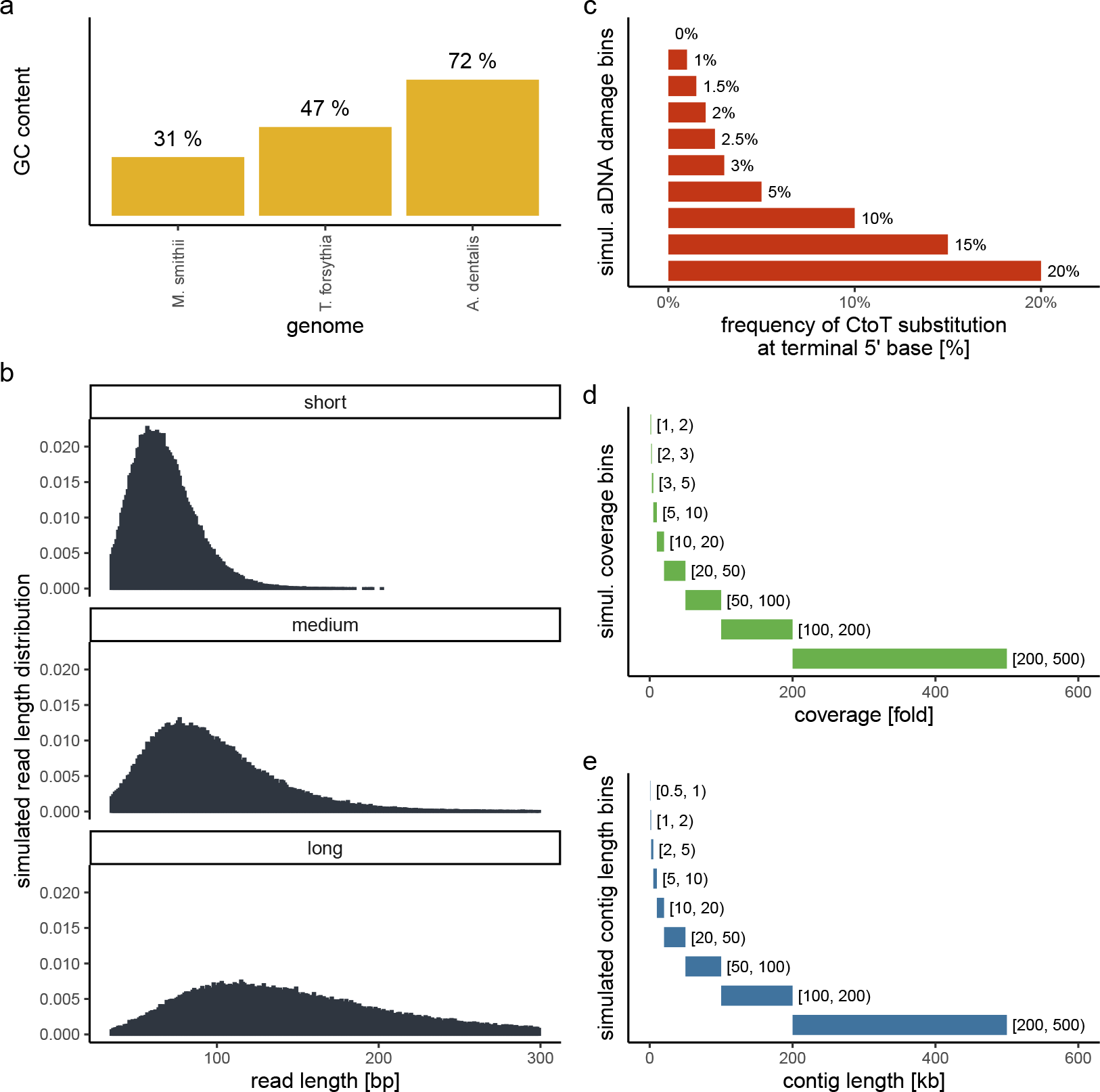
Simulation scheme for evaluating the performance of PyDamage. **a** The GC content of the three microbial reference genomes. **b** The read length distributions used as input into gargammel *fragSim*. **c** The amount of aDNA damage as observed as the frequency of C-to-T substitutions on the terminal 5’ end of the DNA fragments that was added using gargammel *deamSim*. **d** Nine coverage bins from which the exact coverage was sampled by randomly drawing a number from the uniform distribution defining the bin. **e** Nine contig length bins from which the exact contig length was sampled by randomly drawing a number from the uniform distribution defining the bin.

Test contigs of different length were simulated by defining nine contig length bins ranging from 0.5 kb to 500 kb length (Figure 1e) and randomly drawing 100 contig lengths from the respective uniform distribution defining each bin. Next, we chose the location of these test contigs by randomly selecting a contig from all contigs of sufficient length. We determined the exact location on the selected test contig from the reference genome by randomly drawing the start position from the uniform distribution defined by the length of the selected reference contig. This resulted in 900 test contigs per reference genome. Using these test contigs, we selected the aligned DNA fragments of the simulated sequencing data that overlapped the region defined by the contig and evaluated them using PyDamage. In total, we evaluated 702,900 test contigs (243,000 contigs for both *M. smithii* and *T. forsythia*, and 216,000 contigs for *A. dentalis*, for which no reference contig longer than 200 kb was available).

### Archaeological sample

#### Preparation and sequencing

We re-analyzed ancient metagenomic data from the archaeological paleofeces sample ZSM028 (Zape 28) dating to ca. 1300 BP from the site of Cueva de los Muertos Chiquitos, in Mexico, previously published in Borry et al. (2020) (ENA run accession codes ERR3678595, ERR3678598, ERR3678602, ERR3678603, and ERR3678613).

#### Bioinformatic processing

The ZSM028 sample was first trimmed to remove adapters, low quality sequences with Q-scores below 20, and short sequences below 30 bp using AdapterRemoval (Schubert et al., 2016) v2.3.1. The reads were *de novo* assembled into contigs using MetaSPAdes Nurk et al. (2017) v3.13.1 using the non-default k-mer lengths 21, 33, and 45. Reads were then mapped back to the contigs with length > 1,000 bp using Bowtie2 (Langmead and Salzberg, 2012), in the `very-sensitive` mode, while allowing up to 1 mismatch in the seeding process. The alignment files were then given as an input to PyDamage v0.50. Contigs passing filtering thresholds were functionally annotated with Prokka v1.14.6 (Seemann, 2014), using the `--metagenome flag`.

#### Contig Taxonomic Profiling

To investigate the taxonomic profile of the contigs that passed the PyDamage filtering, we ran Kraken2 v2.1.1 (Wood et al., 2019) using the PlusPFP database (https://benlangmead.github.io/aws-indexes/k2) from 27/1/2021. We then generated the Sankey plot using Pavian (Breitwieser and Salzberg, 2016).

### PyDamage implementation

PyDamage takes alignment files of reads (in `SAM`, `BAM`, or `CRAM` format) mapped against reference sequences (*i.e*., contigs, a MAG, a genome, or any other reference sequences of DNA). For each read mapping to each reference sequence *j*, using pysam (pysam developers, 2018), we count the number of apparent C-to-T transitions at each position which is *i* bases from the 5’ terminal end, *i* ∈ {0,1, ⋯, *k*}, denoted 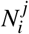 (by default, we set *k*=35). Similarly we denote the number of observed conserved ‘C-to-C’ sites 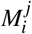, thus

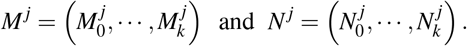

Finally, we calculate the proportion of C-to-T transitions occurring at each position, denoted 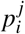 in the following way:

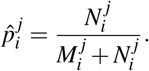

For *D_i_*, the event that we observe a C-to-T transition *i* bases from the terminal end, we define two models: a null model 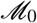(equation 1) which assumes that damage is independent of the position from the 5’ terminal end, and a damage model 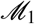 (equation 2) which assumes a decreasing probability of damage the further a the position from the 5’ terminal end. For the damage model, we re-scale the curve to the interval defined by parameters 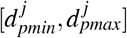.

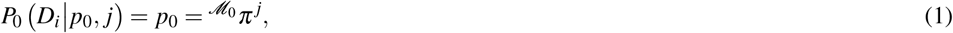

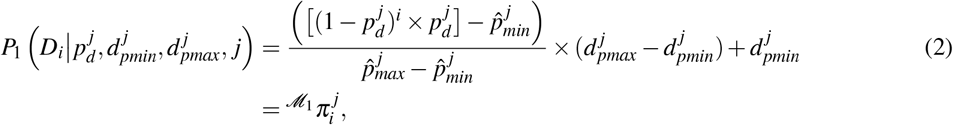

where

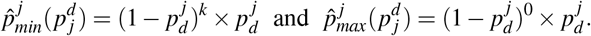

Using the curve fitting function of Scipy (Virtanen et al., 2020), with a `trf` (Branch et al., 1999) optimization and a Huber loss (Huber, 1992), we optimize the parameters of both models using 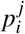, by minimising the sum of squares, giving us the optimized set of parameters

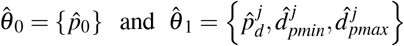

for 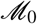 and 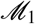 respectively. Under 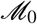 and 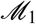 we have the following likelihood functions

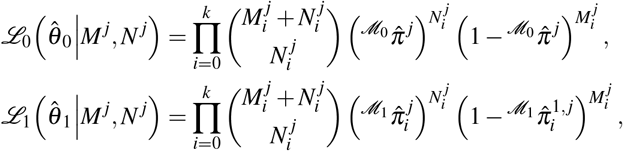

where 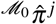 and 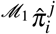 are calculated using equations 1 and 2. Note that if 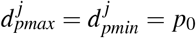, then 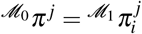 for *i* = 0, ⋯, *k*. Hence to compare the goodness-of-fit for models 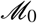 and 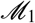 for each reference, we calculate a likelihood-ratio test-statistic of the form

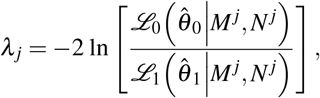

from which we compute a *p*-value using the fact that 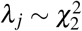, asymptotically. Finally, we adjust the *p*-values for multiple testing of all references, using the StatsModels (Seabold and Perktold, 2010) implementation of the Benjamini-Hochberg procedure (Benjamini and Hochberg, 1995).

## RESULTS

### Statistical Analysis and Model Selection

To test the performance of PyDamage in recognizing metagenome-assembled contigs with ancient DNA damage, we used the simulated short-read sequencing data aligned against simulated contigs of different lengths. Our method correctly identified contigs as not significantly damaged for simulations with no damage in 100% of cases. However, our model only correctly identified contigs as significantly damaged in 87.71% of cases where the contigs were simulated to have damage. To assess the performance of our method, and to determine the simulation parameters that most affected model accuracy, we analysed the simulated data using logistic regression via the `glm` function as implemented in the stats package using R (R Core Team, 2018). We included as potential explanatory variables the median read length, the simulated coverage, the simulated contig length, the simulated level of damage, and the GC content of each of the reference contigs, yielding 32 candidate logistic regression models.

We separated the data into two data sets: half of our data was used as ‘fit data’, data for performing model fit and parameter estimation, and the remaining half was reserved as ‘test data’, data that is used to assess model accuracy on data not used in fitting the model (*n* = 206, 831 in both cases). Unfortunately, with so many observations in our model, classical model selection methods such as AIC and ANOVA tend to overfit (Babyak, 2004). Instead, for each of the fitted 32 logistic regression models (with *ε* = 1 × 10^−14^ and maximum iterations 10^3^) we calculated the ‘balanced accuracy’ (the average of the sensitivity and the specificity) and Nagelkerke’s *R*^2^. We chose the balanced accuracy to equally weight the importance of detecting true damage when it is present, and to also reject a false identification of damage when it is not present.

Of the 32 candidate models, four models had the highest *R*^2^ between 0.556 and 0.568 (compared to the next greatest of 0.429, see Table 1). Of these four models, the maximum balanced accuracy belonged to the model which had the following predictor variables: contig length, mean coverage, GC content and the simulated level of damage, although these values were extremely close for all four models (see Figure 2). Because it is possible that there is correlation between some of our predictor variables (*i.e*. increased levels of simulated damage could lead to a reduced median read length), we then performed a Relative Weights Analysis (RWA) to further estimate predictor variable importance in an uncorrelated setting (Chan, 2020). In essence, RWA calculates the proportion of the overall *R*^2^ for the model that can be attributed to each variable. We performed RWA on both the full model and our best performing model. We found that the median read length and GC content accounted for only 0.31% and 2.75% of the *R*^2^ value in the full model respectively. However, we found that contig length, mean coverage and the simulated level of damage all accounted for approximately one third of the *R*^2^ value in our best performing model, indicating that these are the predictor variables of importance.

**Figure 2.**
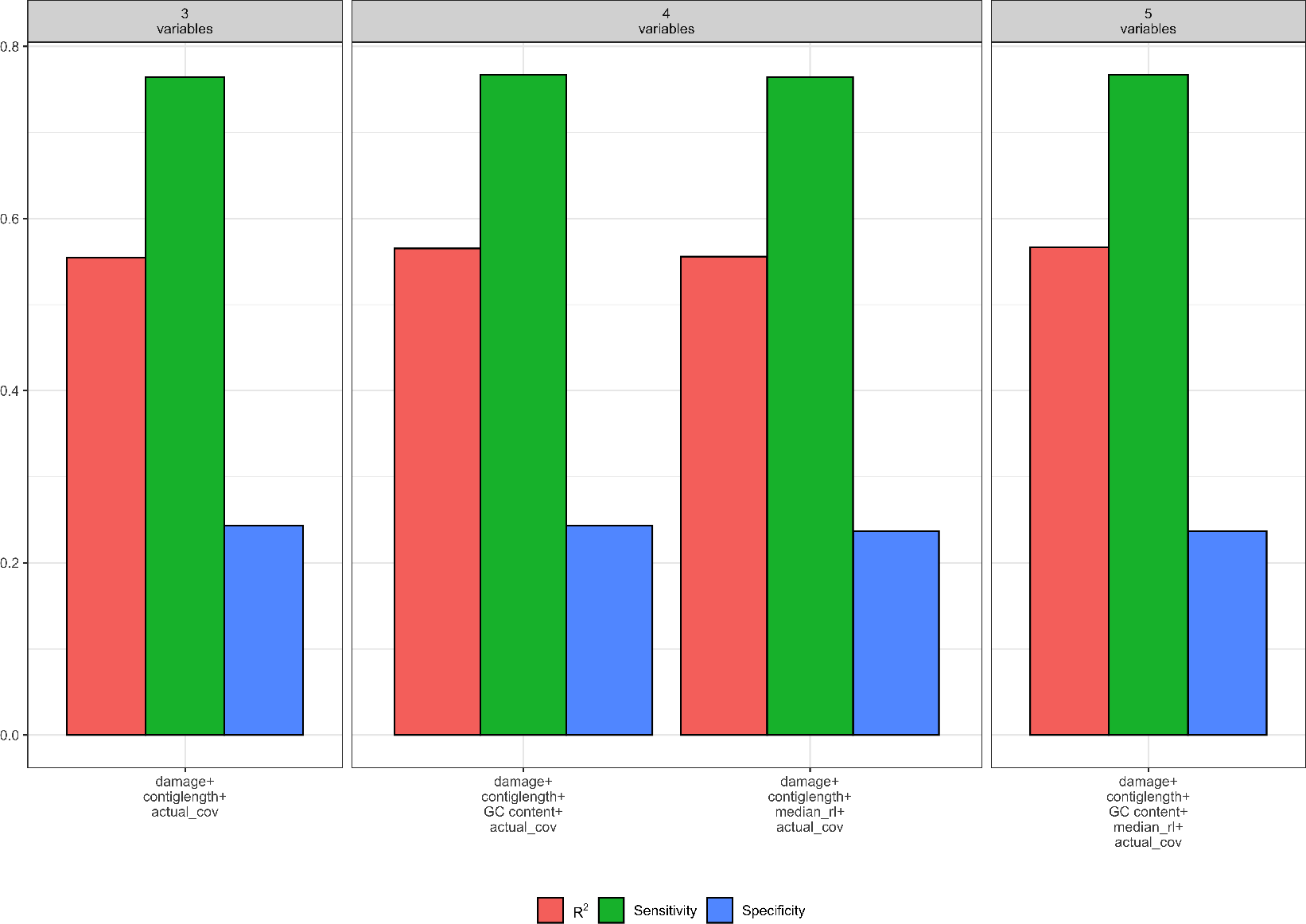
Measures of model fit calculated on the test data for three variables, four variables, and five variables, where red is Nagelkerke’s *R^2^*, green is model sensitivity and blue is model specificity. Model fit is not improved with the addition of variables beyond DNA damage, contig length, and contig coverage.

**Table 1.** Balanced accuracy and Nagelkerke’s *R*^2^ values for the top ten models (as measured by *R*^2^). The model we retained is highlighted in bold.

| Variables | Balanced Accuracy | $R^2$ |
| --- | --- | --- |
| damage/contiglength/GCcontent/readlength/coverage | 0.495 | 0.568 |
| damage/contiglength/GCcontent/coverage | 0.496 | 0.566 |
| damage/contiglength/readlength/coverage | 0.492 | 0.557 |
| **damage/contiglength/coverage** | **0.493** | **0.556** |
| contiglength/GCcontent/readlength/coverage | 0.429 | 0.429 |
| contiglength/GCcontent/coverage | 0.430 | 0.428 |
| contiglength/readlength/coverage | 0.421 | 0.420 |
| contiglength/coverage | 0.422 | 0.419 |
| damage/contiglength/GCcontent/readlength | 0.574 | 0.378 |
| damage/contiglength/GCcontent | 0.581 | 0.377 |

Our final logistic regression model identified mean coverage, the level of damage, and the contig length as significant predictor variables for model accuracy. Each of these variables had positive coefficients, meaning that an increase in damage, genome coverage, or contig length all lead to improved model accuracy. Each variable contributed about one third weight to the *R*^2^ value in the model, indicating roughly equal importance in the accuracy of PyDamage. We integrated the best logistic regression model in PyDamage, with the StatsModels (Seabold and Perktold, 2010) implementation of GLM to provide an estimation of PyDamage ancient contig prediction accuracy given the amount of damage, coverage, and length for each reference (Figure 3), and found these predictions to adequately match the observed model accuracy for our simulated data set (Figure 4).

**Figure 3.**
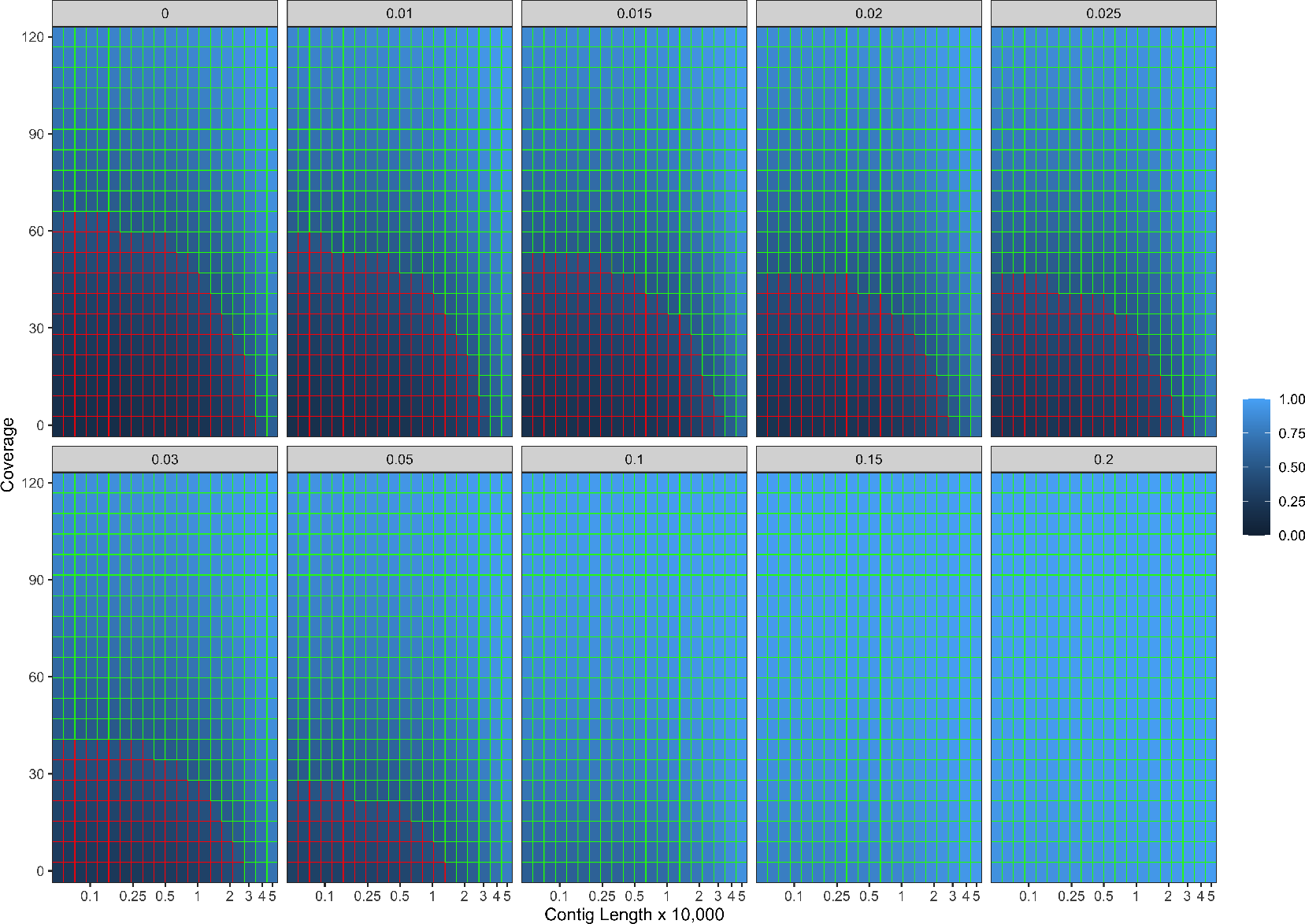
Predicted model accuracy of simulated data. Light blue indicates improved model accuracy, with parameter combinations resulting in better than 50% accuracy are outlined in green.

**Figure 4.**
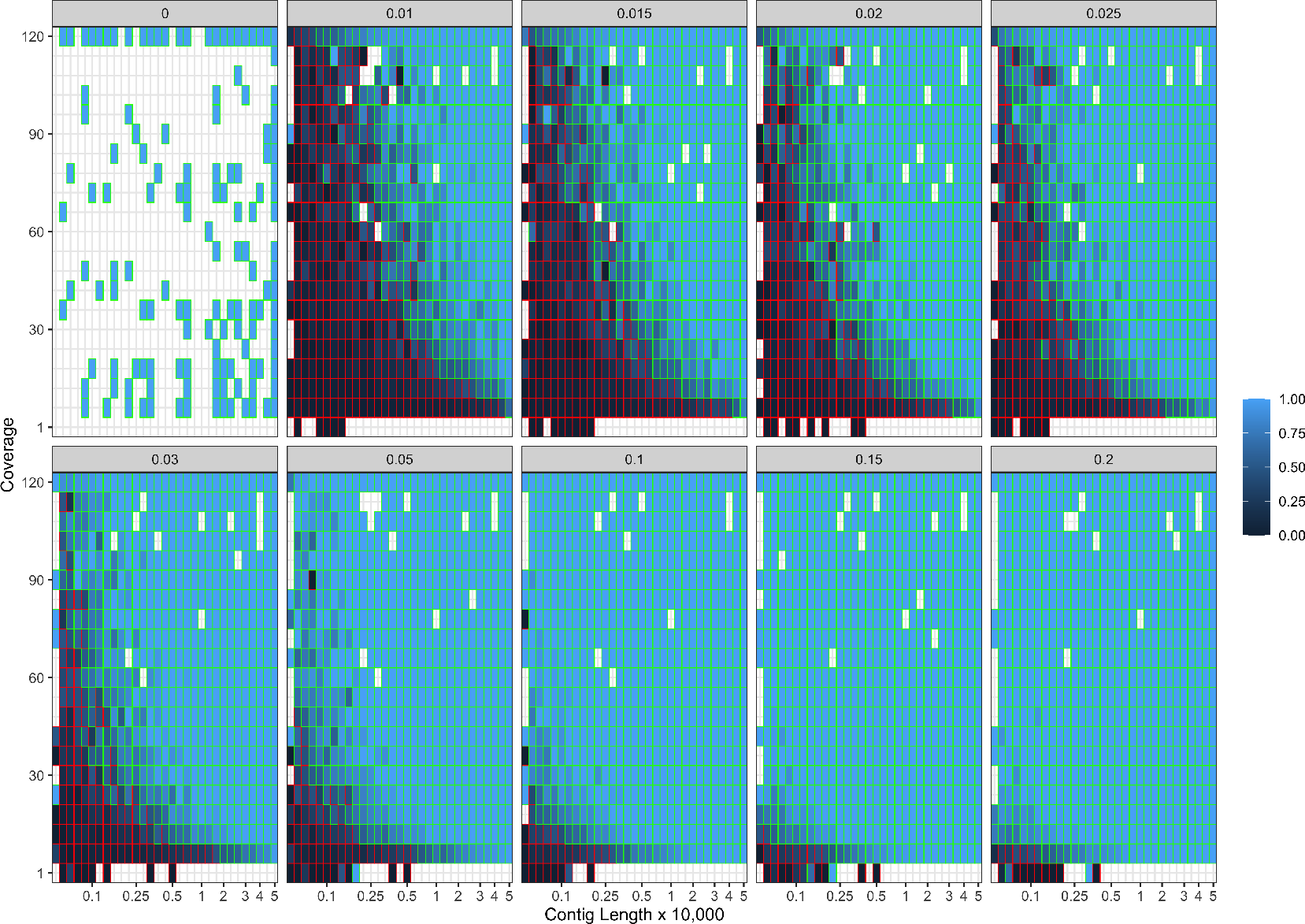
Observed model accuracy of simulated data. Light blue indicates improved model accuracy, with parameter combinations resulting in better than 50% accuracy are outlined with green lines. Grey tiles represent parameter combinations that were not sampled.

### Application of PyDamage to Archeological samples

To test PyDamage on empirical data, we assembled metagenomic data from the paleofeces sample ZSM028 with the metaSPAdes *de novo* assembler. We obtained a total of 359,807 contigs, with an N50 of 429 bp. Such assemblies, consisting of a large number of relatively short contigs, are typical for *de novo* assembled aDNA datasets (Wibowo et al., 2021). After filtering for sequences longer than 1,000 bp, 17,103 contigs were left. PyDamage was able to perform a successful damage estimation for 99.75% of these contigs (17,061 contigs). Because the ZSM028 sequencing library was not treated with uracil-DNA-glycosylase (Rohland et al., 2015), nor amplified with a damage suppressing DNA polymerase, we expect a relatively shallow DNA damage decay curve, and thus filtered for this using the 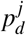 parameter. We chose a prediction accuracy threshold of 0.67 after locating the knee point on Figure 5 with the kneedle method (Satopaa et al., 2011). After filtering PyDamage results with a *q*-value≤ 0.05, 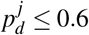, and *prediction accuracy* ≥ 0.67, 1,944 contigs remain. The 5’ damage for these contigs ranges from 4.0% to 45.1% with a mean of 14.3% (Figure 7). Their coverage spans 6.1*X* to 1,579.8*X* with a mean of 65.6*X*, while their length ranges from 1,002 bp to 90,306 bp with a mean of 5,212 bp and an N50 of 10,805 bp.

**Figure 5.**
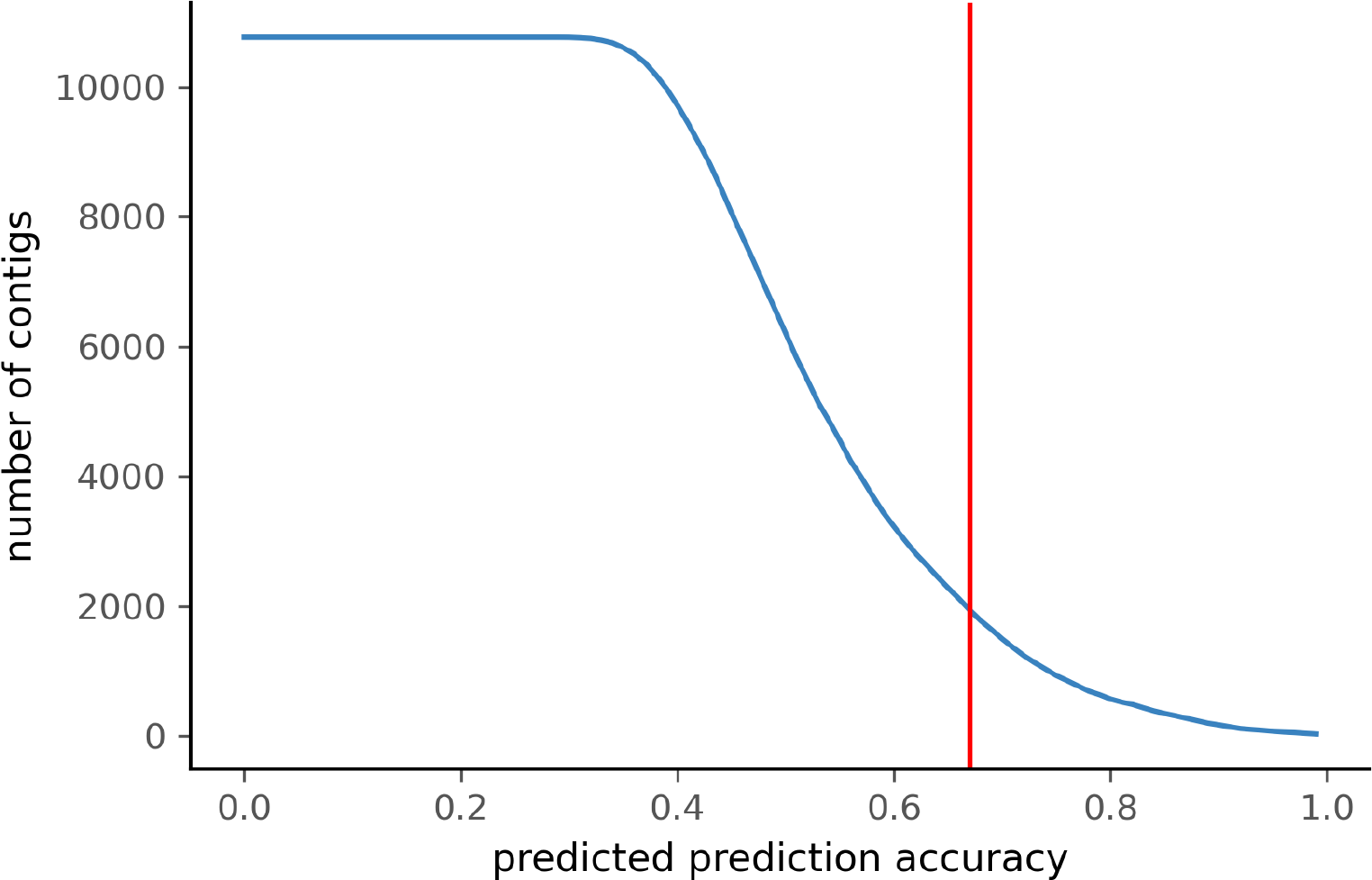
Number of ZSM028 contigs filtered by PyDamage with a *q*-value ≤ 0.05 as a function of the predicted prediction accuracy. In total, 12,271 of the 17,061 contigs were assigned *q*-value ≤ 0.05.

The Kraken2 taxonomic profile of the microbial contigs identified by PyDamage identified as ancient (Figure 6) is consistent with bacteria known to be members of the human gut microbiome, including *Prevotella* (239 contigs), *Treponema* (166 contigs), *Bacteroides* (103 contigs), *Lachnospiraceae* (119 contigs) *Blautia* (36 contigs), *Ruminococcus* (25 contigs), *Phocaeicola* (18 contigs) and *Romboutsia* (16 contigs) (Schnorr et al., 2016; Pasolli et al., 2019; Singh et al., 2017), as well as taxonomic groups known to be involved in initial decomposition, such as *Clostridium* (145 contigs) (Hyde et al., 2017; Harrison et al., 2020; Dash and Das, 2020). In addition, eukaryotic contigs were assigned to humans (18 contigs), and to the plant families Fabaceae (18 contigs) and Solanaceae (18 contigs), two families of economically important crops in the Americas that include beans, tomatoes, chile peppers, and tobacco. The remaining contigs were almost entirely assigned to higher taxonomic levels within the important gut microbiome phyla Bacteriodetes, Firmicutes, Proteobacteria, and Spirochaetes, as well as to the Streptophyta phylum of vascular plants. Collectively, these 5 phyla accounted for 1283 of to 1494 contigs that could be taxonomically assigned.

**Figure 6.**
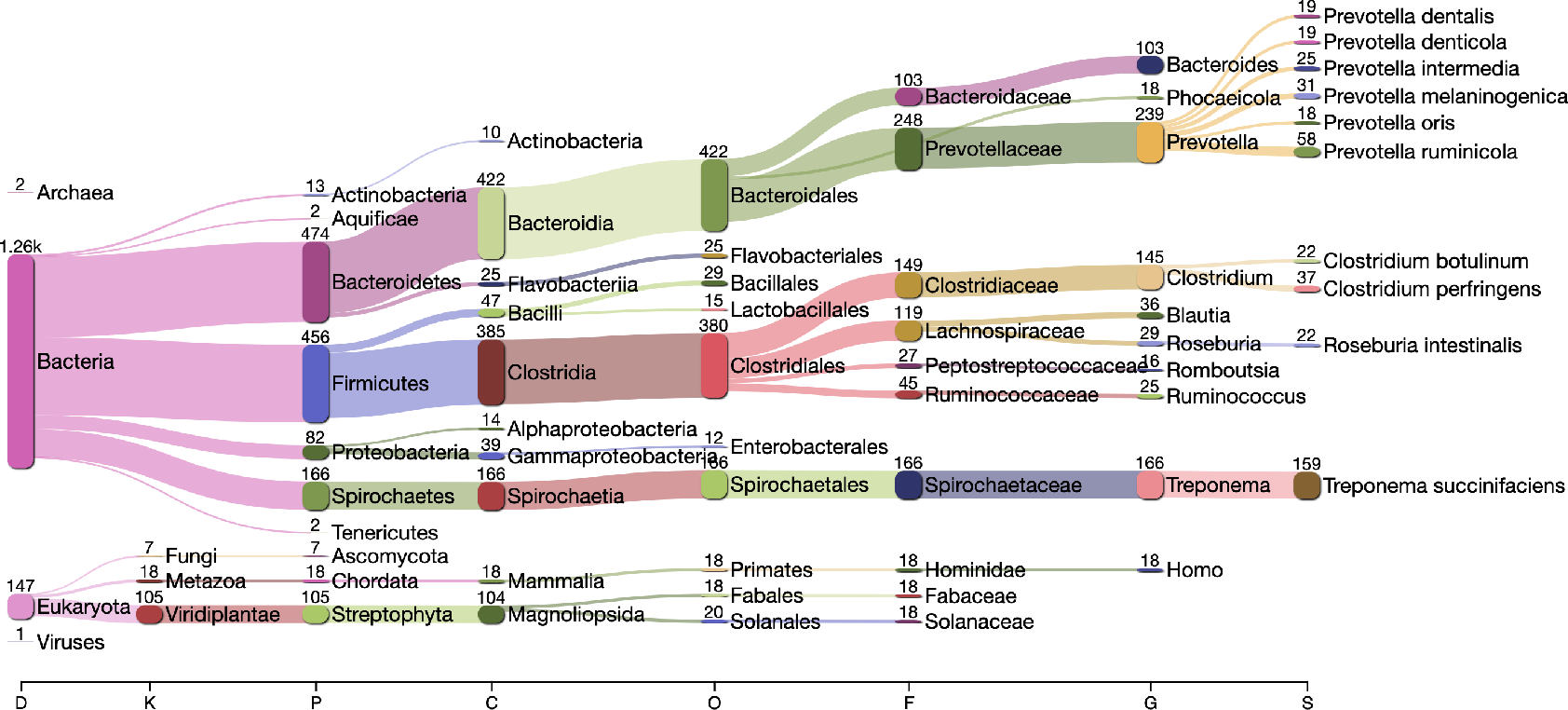
Taxonomic assignation by Kraken2 of the contigs filtered by PyDamage with *q*-value≤ 0.05, 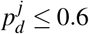, and *prediction accuracy* ≥ 0.67

**Figure 7.**
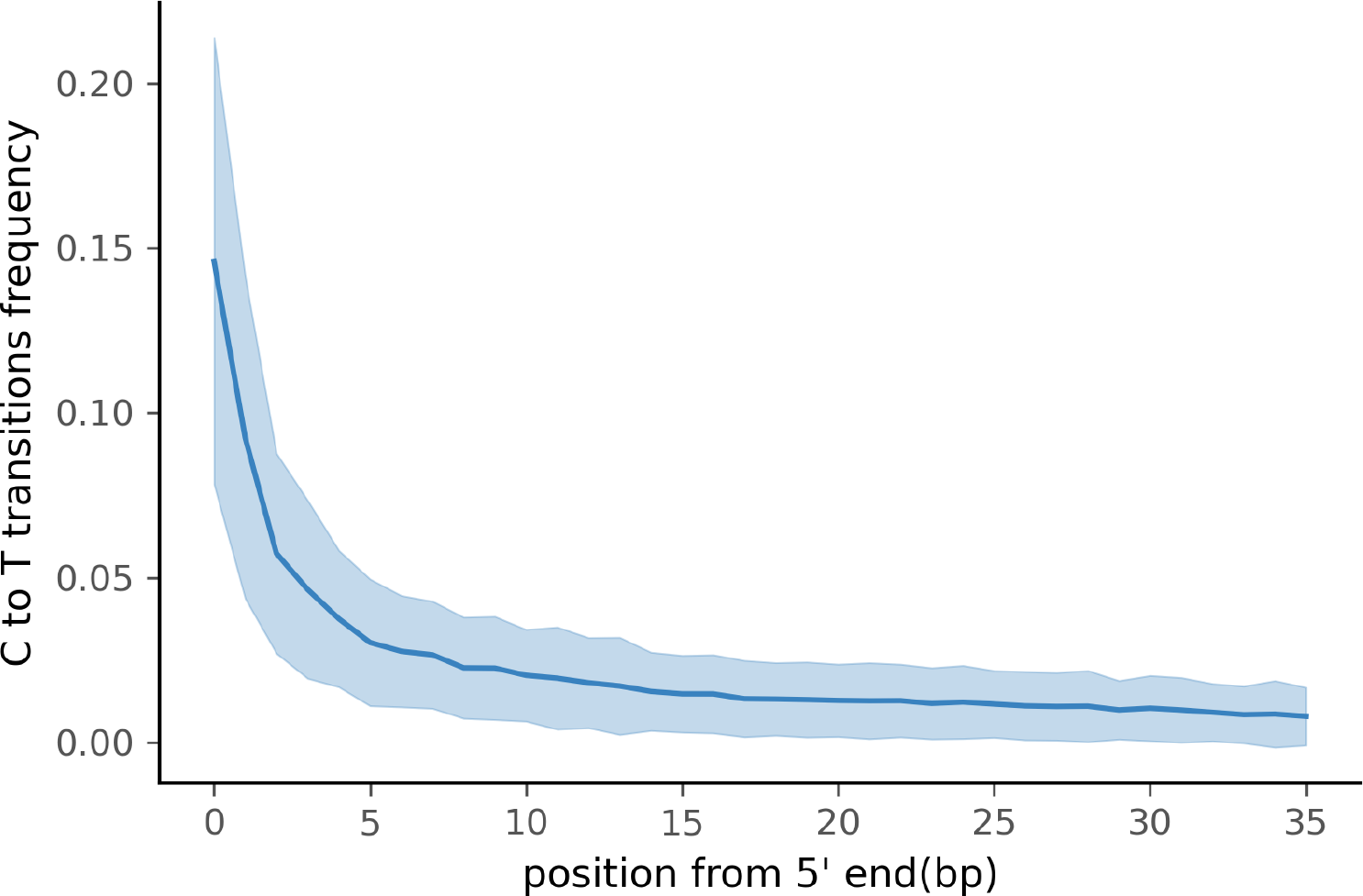
Damage profile of PyDamage filtered contigs of ZSM028. The center line is the mean, the shaded area is ± one standard-deviation around the mean

Functional annotation of the authenticated ancient contigs using Prokka was successful for 1,901 of 1,944 contigs. Among these, multiple genes of functional interest were identified, including contigs annotated as encoding the multidrug resistance proteins MdtA, MdtB, and MdtC, which convey, among other functions, bile salt resistance (Nagakubo et al., 2002) (Table 2). Kraken2 taxonomic profiling of these three contigs yields a taxonomic assignation to the gut spirochaete *Treponema succinifaciens*, a species absent in the gut microbiome of industrialized populations, but which is found globally in societies practicing traditional forms of subsistence (Obregon-Tito et al., 2015; Schnorr et al., 2014). Other authenticated contigs contained genes associated with resistance to the natural antimicrobial compounds fosmidomycin, colistin, daunorubicin/doxorubicin, tetracycline, polymyxin, and linearmycin. A growing body of evidence supports an ancient origin for resistance to most classes of natural antibiotics (D’Costa et al., 2011; Warinner et al., 2014; Christaki et al., 2020; Wibowo et al., 2021).

**Table 2.** Contigs assembled by metaSPAdes, identified by PyDamage as carrying damage, and annotated as carrying resistance genes by Prokka

| contig name | contig length (bp) | coverage | product |
| --- | --- | --- | --- |
| NODE_2446 | 3232 | 64.3 | Arsenical-resistance protein Acr3 |
| NODE_45 | 28638 | 26.0 | Bifunctional polymyxin resistance protein ArnA |
| NODE_832 | 6259 | 46.3 | Cobalt-zinc-cadmium resistance protein CzcA |
| NODE_832 | 6259 | 46.3 | Cobalt-zinc-cadmium resistance protein CzcB |
| NODE_2661 | 3058 | 91.5 | Colistin resistance protein EmrA |
| NODE_2661 | 3058 | 91.5 | Colistin resistance protein EmrA |
| NODE_215 | 13020 | 27.0 | Daunorubicin/doxorubicin resistance ATP-binding protein DrrA |
| NODE_136 | 16294 | 26.0 | Daunorubicin/doxorubicin resistance ATP-binding protein DrrA |
| NODE_1676 | 4090 | 81.3 | Fosmidomycin resistance protein |
| NODE_8410 | 1542 | 77.3 | Linearmycin resistance ATP-binding protein LnrL |
| NODE_29 | 35207 | 27.8 | Multidrug resistance ABC transporter ATP-binding and permease protein |
| NODE_232 | 12485 | 31.9 | Multidrug resistance protein MdtA |
| NODE_97 | 19553 | 27.4 | Multidrug resistance protein MdtA |
| NODE_12 | 45672 | 45.6 | Multidrug resistance protein MdtA |
| NODE_10 | 46280 | 59.8 | Multidrug resistance protein MdtA |
| NODE_97 | 19553 | 27.4 | Multidrug resistance protein MdtB |
| NODE_97 | 19553 | 27.4 | Multidrug resistance protein MdtB |
| NODE_12 | 45672 | 45.6 | Multidrug resistance protein MdtC |
| NODE_10 | 46280 | 59.8 | Multidrug resistance protein MdtC |
| NODE_232 | 12485 | 31.9 | Multidrug resistance protein MdtC |
| NODE_17 | 41269 | 29.9 | Multidrug resistance protein MdtK |
| NODE_465 | 8695 | 37.5 | Tetracycline resistance protein TetO |
| NODE_204 | 13262 | 44.9 | Tetracycline resistance protein, class C |

## DISCUSSION AND CONCLUSION

*De novo* sequence assembly is increasingly being applied to ancient metagenomic data in order to improve lower rank taxonomic assignment and to enable functional profiling of ancient bacterial communities. The ability to reconstruct reference-free ancient genes, gene complexes, or even genomes opens the door to exploring microbial evolutionary histories and past functional diversity that may be underrepresented or absent in present-day microbial communities. A critical step in reconstructing this past diversity, however, is being able to distinguish DNA of ancient and modern origin (Warinner et al., 2017). Characteristic forms of damage that accumulate in DNA over time, such as DNA fragmentation and cytosine deamination, are widely used to authenticate aDNA (Orlando et al., 2021) and have been important, for example, in enabling the reconstruction of the Neanderthal genome from skeletal remains contaminated with varying levels of modern human DNA (Briggs et al., 2007a; Bokelmann et al., 2019; Peyrégne et al., 2019).

Nevertheless, applying such an approach to complex ancient microbial communities, such as archaeological microbiome samples or sediments, is more challenging. Existing microbial reference sequences in databases such as NCBI RefSeq have been found to be insufficiently representative of modern microbial diversity (Pasolli et al., 2019; Manara et al., 2019), let alone ancient diversity, making reference-free *de novo* assembly particularly desirable for both modern and ancient microbial metagenomics. However, *de novo* assembly of aDNA has always been a challenge due to its highly fragmented nature. While tools have been designed to improve the assembly of ancient metagenomics data (Seitz and Nieselt, 2017), assessing the damage carried by the assembled contigs has remained an open problem. Although existing tools can determine the degree of aDNA damage for sequences mapped to a given reference sequence, scaling this up to accommodate the tens to hundreds of thousands of contig references generated by metagenomics assembly requires an alternative, automated approach to damage estimation.

Here, we have presented PyDamage as a tool to rapidly assess aDNA damage patterns for numerous reference sequences in parallel, allowing damage profiling of metagenome assembled contigs. To evaluate the performance of PyDamage model fitting and statistical testing, we benchmarked the tool using simulated assembly data of known coverage, length, GC content, read length, and damage level. While GC content and read length were not a major driver of the accuracy of PyDamage’s predictions, reference length, coverage, and damage level each played major roles. Taken together, this three parameter combination greatly influenced the ability of PyDamage to make a accurate damage assessments for a given contig. Overall, PyDamage has highly reliable damage prediction accuracy for contigs with high coverage, long lengths, and high damage, but the tool’s power to assess damage is reduced for lower coverage, shorter contigs length, and lower deamination damaged contigs. Although aDNA damage levels (cytosine deamination and fragmentation) are features of the DNA itself and out of the researcher’s control, we show that researchers can generally improve model accuracy through deeper sequencing.

When comparing the parameter range of our simulated data to real world *de novo* assembly data, we find that some of PyDamage prediction accuracy limitations are mitigated by the assembly process itself: *de novo* assemblers usually need a minimum of approximately 5X coverage to assemble contigs (Figure 8) (Wibowo et al., 2021), and it is common practice to discard short contigs (<1000 bp) before further processing steps in a classical metagenomic *de novo* assembly analysis process. Nevertheless, low coverage, low damage, short contigs will remain a marginal challenge for damage characterization, even with further manual inspection. For example, for a 10,000 bp *de novo* assembled contig with 10% damage, PyDamage will only start to make reliable predictions once a coverage of 16X is reached (Figure 3). For a similar contig with 20% damage, model accuracy is high even from 1X coverage. Overall, we find that PyDamage generally performs well on ancient metagenomic data with >5% damage, but contig length and coverage are also essential factors in determining the model accuracy for a given contig.

**Figure 8.**
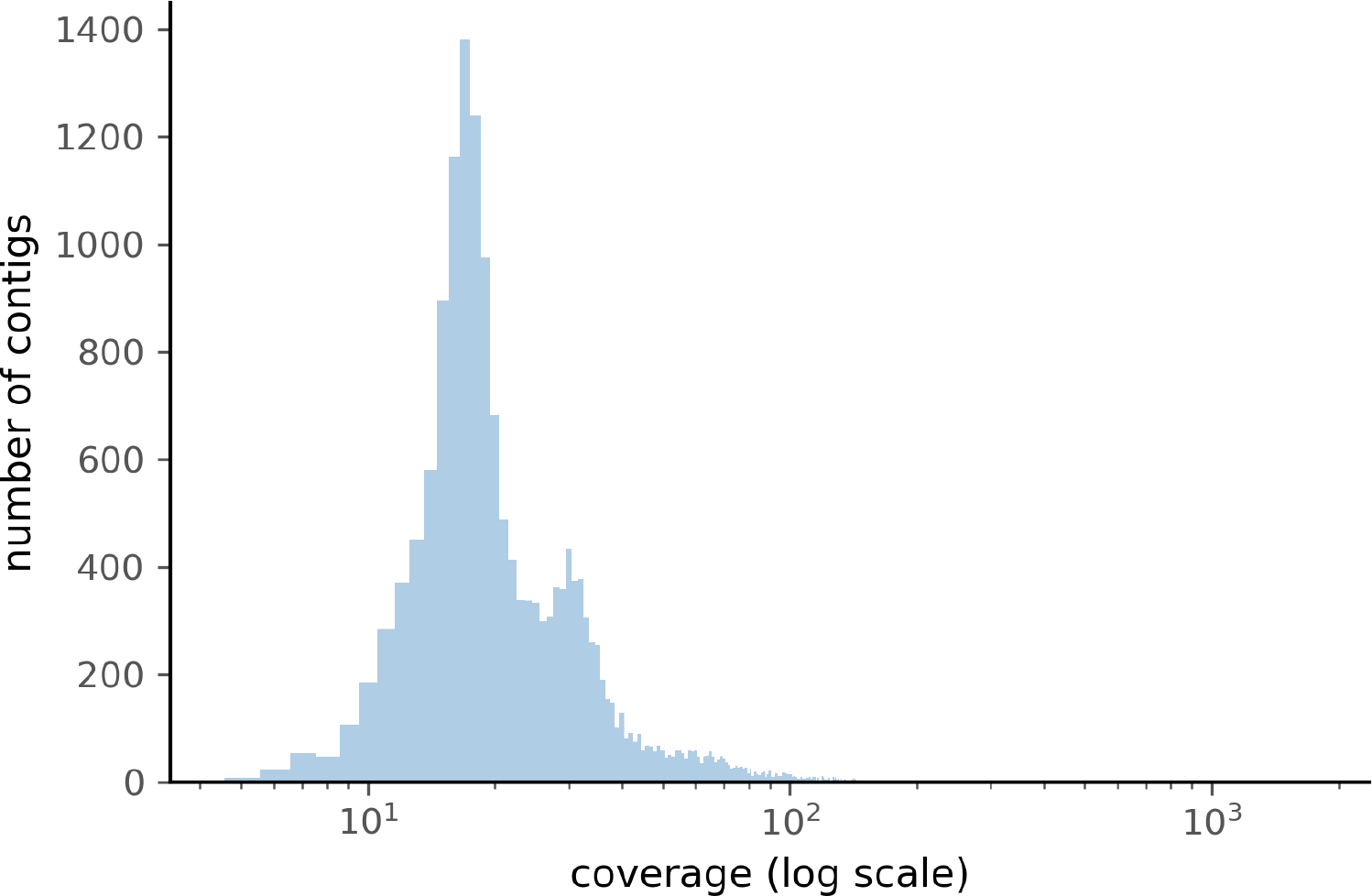
Distribution of the coverage for ZSM028 contigs > 1,000 bp assembled by metaSPAdes.

Although we used the kneedle method (Satopaa et al., 2011) to select the prediction accuracy threshold for paleofeces sample ZSM028, users can adjust the selected prediction accuracy threshold according to the needs of their research question. For example, for some research questions where high accuracy in verifying damage is paramount, more stringent thresholds can be applied to minimize false positives, even though this increases false negatives. For other questions and where additional authentication criteria are available (such as taxonomic information or metagenomic bins), lower thresholds may be applied to reduce the number of false negatives due to insufficient coverage or contig length.

PyDamage is designed to estimate accumulated DNA damage in *de novo* assembled metagenomic sequences. However, although DNA damage can be used to authenticate DNA as ancient, it is important to note that it is not necessarily an indicator of *intra vitam* endogeneity. DNA within ancient remains typically consists of both an endogenous fraction present during life and an exogenous fraction accumulated after death. For skeletal remains, the endogenous fraction typically consists of host DNA, as well as possibly pathogen DNA if the host was infected at the time of death. For paleofeces or dental calculus, the endogenous fraction typically consists of microbiome DNA, as well as trace amounts of host, parasite, and dietary DNA. In both cases, the endogenous fraction of DNA is expected to carry DNA damage accumulated since the death (skeletal remains, dental calculus) or defecation (paleofeces) of the individual. Within the exogenous fraction, however, the DNA may span a range of ages. Nearly all ancient remains undergo some degree of degradation and decomposition, during which either endogenous (thanatomicrobiome) or exogenous (necrobiome) bacteria invade the remains and grow (Hyde et al., 2017; Harrison et al., 2020; Dash and Das, 2020). DNA from bacteria that participated early in this process (shortly after death or defecation), will carry similar levels of damage as the endogenous DNA because they are of similar age. In contrast, more recent necrobiome activity will carry progressively less age-related damage, and very recent sources of contamination from excavation, storage, curation, and laboratory handling are expected to carry little to no DNA damage.

To demonstrate the utility of PyDamage on ancient metagenomic data, we applied PyDamage to paleofeces ZSM028, a ca. 1300-year-old specimen of feces from a dry rockshelter site in Mexico that was previously shown to have excellent preservation of endogenous gut microbiome DNA and low levels of environmental contamination (Borry et al., 2020). Using PyDamage, we assessed the damage profiles of contigs with lengths >1,000 bp, and authenticated nearly 2,000 contigs as carrying damage patterns consistent with ancient DNA. The overwhelming majority of these contigs were consistent with bacterial members of the human gut microbiome, as well as expected host and dietary components, but a small fraction of authenticated contigs were assigned to environmental bacteria and fungi, including the exogenous soil bacteria *Clostridium botulinum* (22 contigs) and *Clostridium perfringens* (38 contigs). These taxa are known to be important early decomposers in the necrobiome (Harrison et al., 2020), and the damage they carry suggests that they likely participated in the early degradation of the paleofeces before decomposition was arrested by the extreme aridity of the rockshelter.

Among the PyDamage authenticated contigs assigned to gut-associated taxa, `NODE_10`, `NODE_12`, and `NODE_97` are of particular interest. These contigs encode a multidrug resistant ABC (MdtABC) transporter associated with bile salt resistance in the bacterium *T. succinifaciens. T. succinifaciens* is a human-associated gut species that is today only found in the gut microbiomes of individuals engaging in traditional forms of dietary subsistence (Obregon-Tito et al., 2015; Schnorr et al., 2014; Angelakis et al., 2019). It is not found in the gut microbiomes of members of industrialized societies, and is believed extinct in these groups (Schnorr et al., 2016). Its identification within paleofeces provides insights into the evolutionary history of this enigmatic microorganism and its functional adaptation to the human gut (Schnorr et al., 2019). The additional identification of other resistance genes among the authenticated contigs provides further evidence regarding the evolution of antimicrobial resistance in human-associated microbes.

As the fields of microbiology and evolutionary biology increasingly turn to the archaeological record to investigate the rich and dynamic evolutionary history of ancient microbial communities, it has become vital to develop tools for assembling and authenticating ancient metagenomic DNA. Coupled with aDNA *de novo* assembly, PyDamage opens up new doors to explore and understand the functional diversity of ancient metagenomes.

## Code and Data availability

- Genetic data for ZSM028 is available on the European Nucleotide Archive (ENA) under accession PRJEB33577.
- PyDamage Software and source code available from: github.com/maxibor/pydamage, license: GPLv3
- The code to replicate the simulation of reads and contigs, and the figures is available in the following citable GitHub repository: DOI: 10.5281/zenodo.4630383 - github.com/maxibor/pydamage-article

## ACKNOWLEDGMENTS

We thank Nigel Bean and Jonathon Tuke for extremely useful discussions. AH was funded by the Deutsche Forschungsgemeinschaft (DFG, German Research Foundation) under Germany’s Excellence Strategy (EXC 2051 – Project-ID 390713860). ABR was funded by the European Research Council (ERC) under the European Union’s Horizon 2020 research and innovation program under grant agreement no. 771234 - PALEoRIDER. MB and CW were funded by the Werner Siemens Foundation (”Paleobiotechnology”).

